# Influenza A viruses serially passaged in different MDCK cell lines exhibit limited sequence variation across their genomes, with the exception of the hemagglutinin gene

**DOI:** 10.1101/2020.02.20.959015

**Authors:** Karen N. Barnard, Brian R. Wasik, Brynn K. Alford-Lawrence, Jessica J. Hayward, Wendy S. Weichert, Ian E.H. Voorhees, Edward C. Holmes, Colin R. Parrish

## Abstract

New methods for deep sequence analysis provide an opportunity to follow the emergence and dynamics of virus mutations in real time. Although viruses are commonly grown in cell culture for research and for vaccine development, the cells used to grow the virus are often not derived from the same tissue or even the same host that the virus naturally replicates in. The selective pressures of culturing virus *in vitro* are still only partially understood. MDCK cells are the standard cell for growing influenza viruses yet are derived from the epithelium of the canine kidney and are also heterogenous. We passaged human H3N2, H1N1 pandemic, and canine H3N2 influenza A viruses (IAV) in different lineages of MDCK cells, as well as lines engineered to express variant Sia receptors, including α2,3- and α2,6-linkages or *N*-glycolylneuraminic acid (Neu5Gc) or *N*-acetylneuraminic acid (Neu5Ac) forms. MDCK-Type II cells had lower infection efficiency and virus production, and infection appeared more dependent on protease activation of the virus. When viruses were passaged in the different cells, they exhibited only small numbers of consensus-level mutations, and most were within the HA gene. Both human IAVs showed selection for single nucleotide minority variants in the HA stem across cell types, as well as low frequency variants in the HA receptor binding site of virus passaged in cells expressing Neu5Gc. Canine H3N2 also showed minority variants near the receptor-binding site in cells expressing Neu5Gc and also in those expressing *α*2,6-linkages.

**IMPORTANCE:** The genetic variation and adaptability of viruses are fundamental properties that allow their evolutionary success in the face of differing host environments and immune responses. The growth of viruses in cell culture is widely used for their study and for preparing vaccines. However, the selection pressures that cell passaging imposes on viruses are often poorly understood. We used deep sequence analysis to define, in detail, how three different influenza A viruses respond to passaging in different lineages of canine MDCK cells that are commonly used for their growth, as well as in variant cells engineered to express different forms of their cell surface receptor, sialic acid. This analysis revealed that most mutations occur in the HA gene and few sequence changes in the virus population reached high proportions. This is relevant for understanding the selective pressures of virus growth in cell culture and how it shapes evolutionary patterns.

## INTRODUCTION

High levels of genetic variation are a characteristic feature of RNA viruses. Such variation fuels evolution in the face of antibody immunity, anti-viral drug treatment, and emergence into new host species, enabling viruses to continue to successfully propagate and transmit over extended periods of time. While the natural and experimental evolution of viruses has been described many times, the underlying processes of sequence variation and natural selection that drive evolution of viral genomic sequences at the scale of individual populations are still not well understood. In this study, we define the detailed dynamics of the sequence variation underlying the evolution of three different influenza viruses in a number of related *in vitro* models, allowing us to better understand key aspects of different selection or genetic drift on virus populations, particularly related to variation in host cells and receptors.

The growth of viruses in culture is widely used to study their phenotypic properties and many “wild type” viruses used in research have been extensively passaged in culture, as have virus strains used in vaccine development. Despite the frequency with which cell culture is used, the selection pressures imposed on the viruses by different cell cultures are still poorly understood. Importantly, the common environment of cell monolayers grown on a solid substrate, overlain with a rich liquid media containing bovine serum, is clearly different from those present in the tissues or mucosal environments that the cells and viruses normally grow in. Viruses are also often grown in cell lines derived from different species and tissues than those which the virus infects during a natural infection. Influenza A viruses (IAV) are segmented, negative-sense RNA viruses that replicate in respiratory tissues in mammals and in the gastrointestinal tracts of many of their avian reservoir hosts (1, 2). However, Madin-Darby Canine Kidney (MDCK) cells have been considered the gold standard cell line for the culturing of IAV since the mid-1970s (3). These cells are described as having been isolated from the kidney of a Cocker Spaniel dog and established in culture, and display features of proximal tubule epithelial cells (4, 5). Their use for growing IAV likely stems, at least in part, from their ability to grow high titers of many avian and human IAV strains, and also their ability to remain attached to the growth substrate in the presence of trypsin when used in infection media to prime the viral hemagglutinin (HA) glycoprotein for infection. Human viruses also appear to show relatively few mutations after passage in MDCK cells, although the selection of HA mutations occurs in some viruses (3, 6). In contrast, growth of human IAV isolates in embryonated chicken eggs, long used for isolation and growth of many viruses, often results in the selection of mutations in HA associated with the adaptation of viruses to the predominant *α*2,3-linkage of the sialic acid (Sia) receptor present, which is often referred to as the “avian receptor” form (7, 8). The HA mutations that arise during egg passaging may alter the antigenic structure of the HA, which can have consequences for viruses used in vaccines (7, 9, 10).

The original MDCK cells, including the MDCK-NBL2 available from the American Type Culture Collection, have been shown to be highly heterogeneous, and many alternate MDCK cell lineages now exist (4). A number of studies have shown that different MDCK clones vary significantly in their properties, and these have been broadly grouped into “Type I” and “Type II” cells (5, 11–14). MDCK-Type I cells are generally sub-cloned from a low passage of the parental MDCK cell line and have a small, flat, spindle-like morphology with strong tight junctions (giving high electrical resistance) and higher density of NA-K proton pumps (11, 12, 15). In comparison, MDCK-Type II cells have been sub-cloned from a high passage of MDCK cells and are large, cuboidal cells that grow in clusters and are characterized as having “leaky” tight junctions (giving low electrical resistance), with fewer NA-K proton pumps, and a slower growth rate (11, 12, 15). The MDCK-Type I and MDCK-Type II cells also differ in their lipid composition and their ability to transport liquid through the monolayer, resulting in hemicysts (blister-like structures) that form under the impervious cell monolayer (12, 14). New approaches to virus isolation and to vaccine development are also dependent on growth in MDCK cells, often in cells with increased levels of *α*2,6-linked Sia (16). Clearly, understanding the effects on virus population variation due to *in vitro* passaging of IAV in different MDCK cell lines will provide both basic and applied information about key virus:host interactions. For example, previous research using a variety of different sub-clones of MDCK showed that Type II-like cells supported a lower replication of IAV compared to Type I-like clones, possibly due to differences in endogenous protease expression (17). However, it is not known whether the different MDCK cell lineages can select for different IAV variants.

A classic paradigm in studies of IAV host range is that IAV tropism is determined by the linkage of Sia to the underlying carbohydrate chain, or glycan, on the surface of cells (1). In this model, the human-adapted IAV strains preferentially bind to α2,6-linked Sia, the predominant linkage found in the upper respiratory tract of humans (18). In contrast most avian IAV strains bind to higher levels to α2,3-linked Sia, the predominant form in the gastrointestinal tract of birds where the virus replicates (19). Pigs have been described as a “mixing vessel” as they express both α2,6- and α2,3-linked Sia in their respiratory tract, allowing for an increased possibility of co-infection by both mammalian and avian IAV strains, which would permit reassortment (20). However, in addition to linkage type there is also considerable variation in Sia structures, including the addition of a variety of chemical modifications (21–23). The most basic Sia form is *N*-acetylneuraminic acid (Neu5Ac), while the hydroxylated form of Neu5Ac, *N*-glycolylneuraminic acid (Neu5Gc) is one of the most common Sia variants. This modification is synthesized by the cytidine 5’-monophosphate-*N*-acetylneuraminic acid hydroxylase (CMAH) that is highly expressed in many mammals, but which has been independently lost in many vertebrate lineages (24). Many natural hosts of IAV express only Neu5Ac but not Neu5Gc due to loss of CMAH function, including humans, “western” breeds of dogs, and birds, while other hosts such as horses and pigs express high levels of Neu5Gc in their respiratory tracts and in other tissues (24–27). While some effects of Neu5Gc have been characterized in functional assays on hemagglutinin (HA) and neuraminidase (NA) (28–30), it remains unclear whether this modified Sia acts as a selective pressure on the virus or influences pathogenesis.

MDCK cells can be used as a model for modifying Sia properties by engineering the enzymes involved in Sia modification or linkages. Plasmid expression of *β*-galactoside *α*2,6-sialyltransferase 1 (ST6GAL1) has been used to produce variant cells termed MDCK-SiaT1 which express higher levels of α2,6-linked Sia compared to wild-type MDCK cells (31). Human viruses passed on standard MDCK cells often show mutations associated with selection by α2,3-linked Sia, while fewer mutations were observed after passage of viruses in MDCK-SiaT1 cells (6, 32, 33). MDCK-SiaT1 have therefore become the preferred cell line for culturing human IAV, particularly for clinical isolates (16). MDCK cells appear to express little or no detectable Neu5Gc, likely due to their origin in a Western breed of dog (3), and therefore can act as a model for Neu5Gc presence by expression of CMAH via transfection with an expression plasmid.

In this study, we passaged three IAV strains (A/California/04/2009 pH1N1, A/Wyoming/3/2003 H3N2, and A/canine/IL/11613/2015 H3N2) in different MDCK cell lines and examined the effects on sequence variation within virus populations. Each virus stock was derived from reverse genetic plasmids and serially passaged on MDCK-NBL2, MDCK-SiaT1, MDCK-CMAH, MDCK-Type I, or MDCK-Type II cells. The virus populations were analyzed in detail using deep sequencing to determine the effect of different cell types and Sia forms on IAV selection *in vitro*.

## RESULTS

### Development of MDCK cell lines

We used the standard ATCC line of MDCK-NBL2 (MDCK-WT) as the parental line to create the different MDCK cell lines for this study. MDCK-WT cells express both α2,3- and α2,6-linked Neu5Ac Sia, and only trace amounts of Neu5Gc (<1%), either due to uptake from the fetal bovine serum in growth media or possibly from residual CMAH activity (Fig. 1) (34). To create MDCK cells with higher levels of the human-like α2,6-linked Sia, MDCK-WT cells were transfected with an expression plasmid carrying the human *ST6GAL1* gene to generate our own stock of MDCK-SiaT1 cells (31). Compared to MDCK-WT, the MDCK-SiaT1 cells showed increased staining in flow cytometry for the *Sambucus nigra* (SNA) lectin which recognizes α2,6-linked Sia (Fig. 1A). MDCK-WT cells were also transfected with an expression plasmid carrying CMAH and sub-cloned to create a cell line that expressed Neu5Gc (MDCK-CMAH). MDCK-CMAH cells expressed ∼40% of their total Sia as Neu5Gc (Fig. 1C) and maintained consistent Neu5Gc expression across cell passages. These levels of Neu5Gc are comparable to those seen in pig respiratory tissues (Supplemental Table 1).

**Figure 1.**
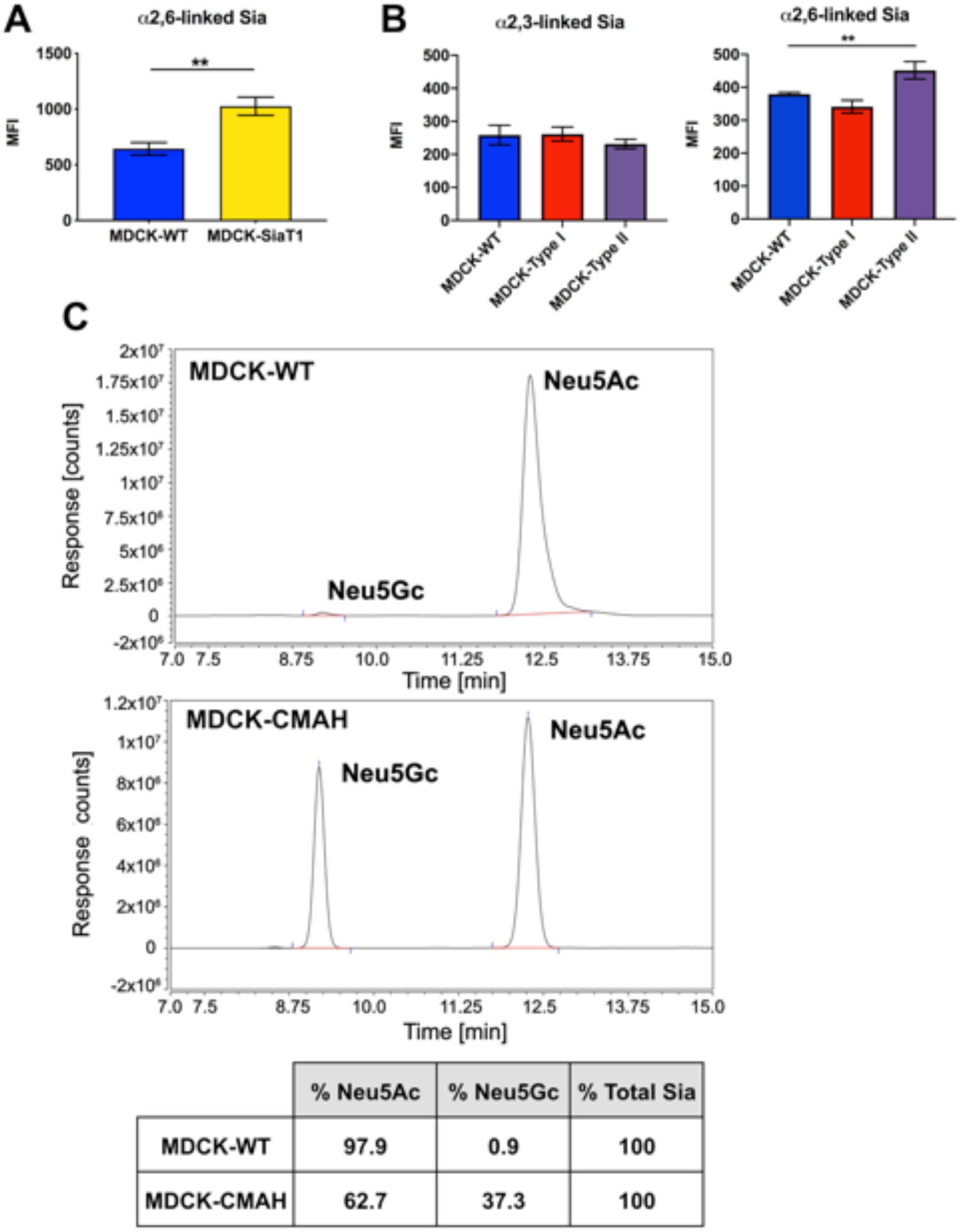
The variation in expression of modified Sia receptors on the different MDCK variants used here. **A)** Plant lectins MAA (α2,3-linked Sia) and SNA (α2,6-linked Sia) were used to compared linkage types between MDCK-WT, MDCK-Type I, and MDCK-Type II cells via flow cytometry. **B)** Plant lectin SNA was used to determine amount of α2,6-linked Sia on MDCK-WT cells compared to MDCK-SiaT1 cells via flow cytometry. **C)** HPLC analysis of total Sia collected from MDCK-WT and MDCK-CMAH cells. Data analyzed by one-way ANOVA using PRISM software. **\*** = p-value ≤0.05; ** = p-value ≤0.01; *** = p-value ≤0.001.

Other MDCK cell variants examined here include clones originally isolated and characterized as described in Nichols et al. (11), which were a kind gift from Dr. William Young (University of Kentucky), and that were defined as MDCK-Type I and MDCK-Type II cell lines. These MDCK-Type I and MDCK-Type II cells expressed the same levels of α2,3-linked Sia as MDCK-WT when stained with *Maakia amurensis type I* (MAA I) lectin, but MDCK-Type II had higher staining for α2,6-linked Sia when stained with SNA lectin (Fig. 1B). The MDCK cell variants were genetically fingerprinted, and all three MDCK cell line samples and all three pairings (MDCK-Type I and MDCK-Type II, MDCK-Type I and MDCK-WT, MDCK-Type II and MDCK-WT) had pi-hat >0.99, suggesting they all originated from the same female Cocker Spaniel-related dog, despite their morphological differences (Table 1).

**Table 1.** Pi-hat measures between each pair of MDCK cell line samples.

|  | MDCK-Type I | MDCK-Type II |
| --- | --- | --- |
| MDCK-Type II | 0.9912 |  |
| MDCK-WT | 0.9992 | 0.9917 |

### Viral passaging and analysis

The study design employed here is diagrammed in Figure 2. To compare results between different IAV strains, we tested three viruses: pandemic H1N1 (pH1N1, A/California/04/2009), a seasonal H3N2 (wyoH3N2, A/Wyoming/3/2003)), and a canine H3N2 strain (CIV H3N2, A/canine/IL/11613/2015). Viruses were recovered from reverse genetics plasmids as previously described (35), and stocks prepared after two passages in MDCK-WT cells. Each virus was titrated by TCID_50_ assay in MDCK-WT cells and passaged five times at low MOI of 0.001 in MDCK-WT, MDCK-CMAH, MDCK-Type I, or MDCK-SiaT1 cells, and MOI of 0.05 in MDCK-Type II cells. Viruses were then passaged for an additional five passages in MDCK-WT, MDCK-CMAH, and MDCK-SiaT1 cells, giving a total of 10 passages. As a second replicate, the same passaging methods were repeated in MDCK-WT and MDCK-CMAH using three separate passage series per cell type for 5 passages to determine the reproducibility of any variation seen. As a positive control for virus selection, we passaged each virus ten times in the presence of increasing concentrations of 4-guanidino-2,4-dideoxy-2,3-dehydro-N-acetylneuraminic acid (Zanamivir), as previously described (36, 37).

**Figure 2.**
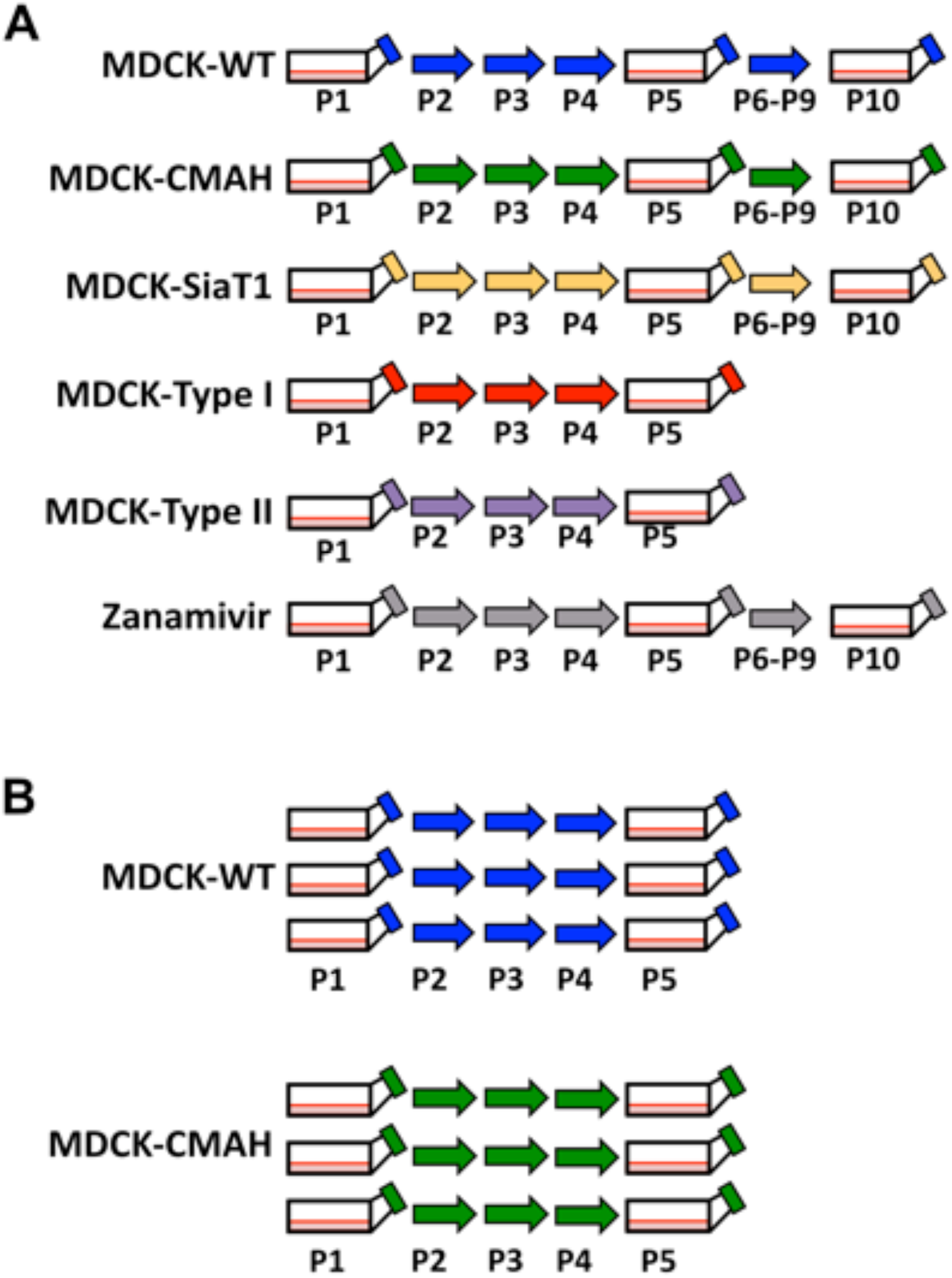
Outline of the design of the experimental evolution studies performed here. **A)** Diagram of replicate 1 for virus passaging in MDCK cells. Passages were completed for human IAV strains wyoH3N2 and pH1N1, along with CIV H3N2 at a low MOI of 0.001. Virus was sequenced for passage 1, 5, and 10. MDCK-Type II cells did not produce enough progeny virus despite increasing MOI, so no virus was sequenced for this cell type. For Zanamivir passaged virus, an MOI of 0.005 was used and Zanamivir concentration was increased during each passage from 0.01 μM to 1 μM. **B)** Replicate 2 was used to determine if there was variation between virus populations passaged in MDCK-WT and MDCK-CMAH, three individual populations of each virus were passaged 5 times. Virus in passage 1 and 5 were sequenced for each population.

Viral RNA was extracted from the stock virus, and from viruses recovered after passage 1, 5, and 10. Viral genomes were amplified using whole genome RT-PCR as previously described (38). Libraries were prepared using Illumina Nextera XT and sequenced using Illumina MiSeq with 2 *×* 250 base paired end reads. To control for error during sequencing, the plasmid stocks for each virus were also deep sequenced (Supplemental Fig. 1). A variant calling threshold of 2% was chosen as we considered that only variants seen above that level at some point in the passaging were likely to be biologically significant. Average coverage for all gene segments for pH1N1 and wyoH3N2 (Supplemental Fig. 2A,B) was 1000 to 10,000 reads per base pair. The coverage for most gene segments for CIV H3N2 were comparably high, with the exception of the HA gene segment that consistently lower coverage (between 50-200 reads per base pair) for reasons that are currently unclear. CIV H3N2 HA-specific primers were therefore used to prepare additional libraries with higher coverage (Supplemental Fig. 2C). The second replicate of the wyoH3N2 passages could not be completed to passage 5 in MDCK-WT cells, as the virus titers dropped below detectable limits during passage 3 for all three flasks. Similarly, CIV H3N2 failed to replicate for all 5 passages in MDCK-Type II cells.

### Passage with Zanamivir

Passaging of virus with the NA inhibitor Zanamivir has previously been shown to select for mutations in both NA and HA as a result of NA inhibition (36, 39, 40). We passaged the three IAVs in increasing concentrations of Zanamivir starting at 0.01 μM in passage 1 up to 50 μM in passage 10. While none of the viruses showed mutations in NA that would affect Sia binding, all three viruses saw single nucleotide variants (SNV) arise in HA near or in the receptor-binding site, although whether these were selected for Zanamivir resistance will require further characterization. For pH1N1 virus, most of these HA SNVs were ≤5% of the population, although V135E reached a frequency of 25% (Fig. 3A). WyoH3N2 gained two SNVs in the HA receptor-binding site that reached high frequencies: A163T (55%) and L244V (47%) (Fig. 3B). In CIV H3N2, the variant S231G in HA reached fixation (Fig. 3C) and was close to a R229I variant that was previously found to confer NA inhibitor resistance in cell-based assays (41). CIV H3N2 also saw fixation of the A27T variant in NA and of the G228R variant in M1 in the M segment; however, both of these variants were also seen in other CIV H3N2 populations, including the stock virus.

**Figure 3.**
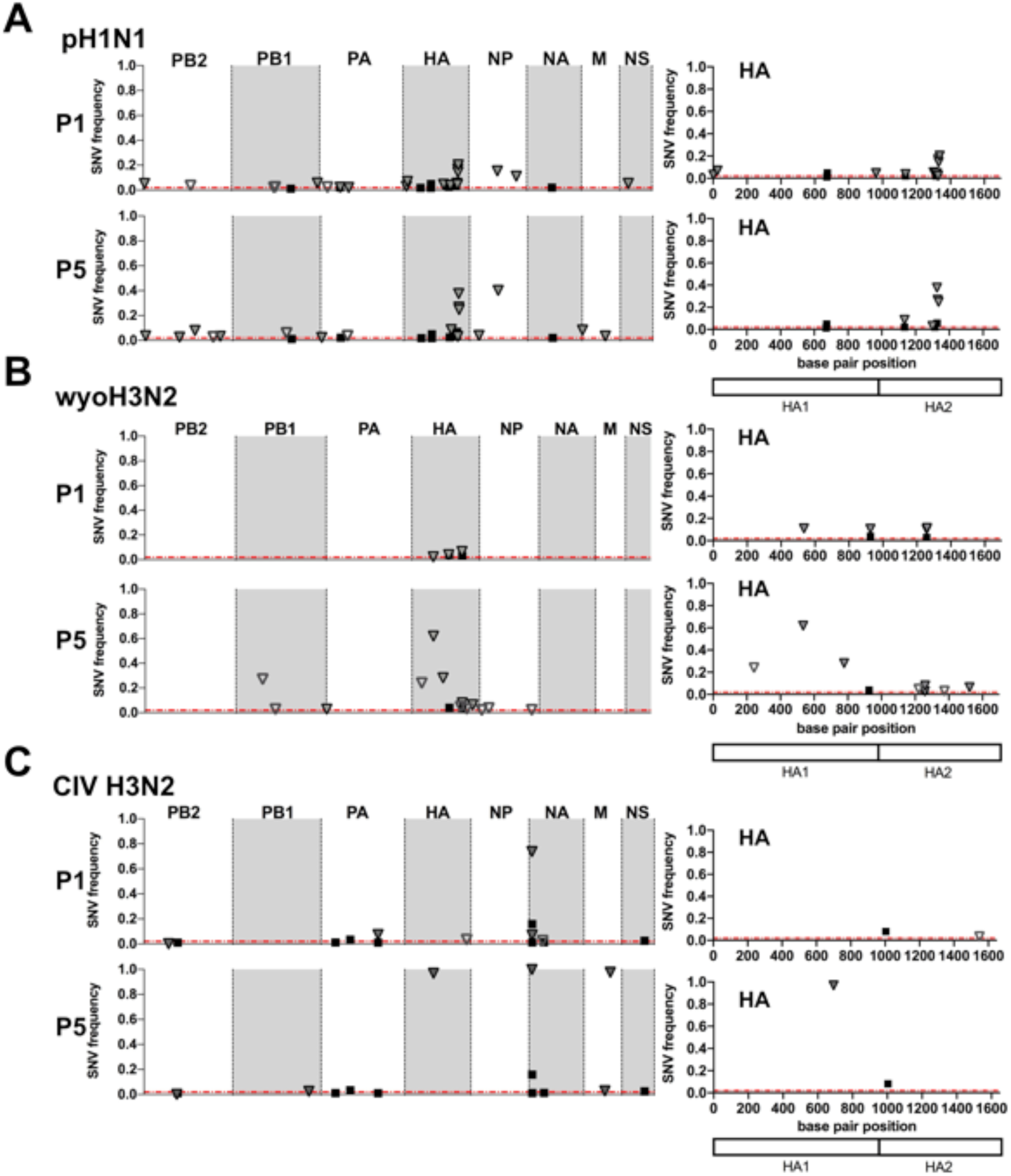
Results of viral passage in the presence of increasing amount of Zanamivir. Single nucleotide variants (SNV) were analyzed for **A)** pH1N1, **B)** wyoH3N2, and **C)** CIV H3N2. Data is shown for the whole genome for passage 1 and passage 5 on the left panel, while the nucleotide position of SNVs for HA are shown on the right side. Non-synonymous mutations are filled in shapes while synonymous mutations are open shapes.

### Passage of pH1N1 in MDCK-WT and MDCK-SiaT1, and MDCK-Type I cells

When pH1N1 was passed in MDCK-WT, MDCK-SiaT1, and MDCK-Type I cells, SNVs were detected in most gene segments; however, most of these were at low frequency, synonymous, and not consistently carried between passages. The HA segment, however, showed a greater number of non-synonymous SNVs and few synonymous changes (Fig. 4; **Supplemental Table 2**). Most SNVs introduced mutations into the HA2 stalk domain, while few SNVs were seen in HA1 corresponding to the receptor binding site domain. None of the SNVs in HA1 were present through passage 5 for MDCK-Type I or passage 10 for MDCK-WT and MDCK-SiaT1 cells. Minority SNVs in the stalk domain increased in frequency between passages 1 and 5 from 25%-45% up to 45%-85%, with the highest frequency being of residue 445, a variant that was also present at low frequency in the stock virus (**Supplemental Table 2**). By passage 10, both MDCK-WT and MDCK-SiaT1 had fixed variants in the stalk domain, with K445E fixed in MDCK-SiaT1 cells and K445M and R509G fixed in MDCK-WT (Fig. 4A,B, **Supplemental Table 2**). MDCK-Type I had a similar stalk variant, T436N, reached a high frequency of 58% (Fig. 4C). Other SNVs clustered in this same region of HA2, including at residues 436, 439, 441, 442, 443, 446, and 448, some at frequencies of 10-50%. Glycosylation sites on HA were maintained across all passaged virus except in one population of MDCK-WT-passaged virus (replicate 2) which lost the proposed glycosylation site at site 21 in passage 1 at a frequency of ∼13%, and maintained this SNV through passage 5 at a frequency of ∼12% (Fig. 5C).

**Figure 4.**
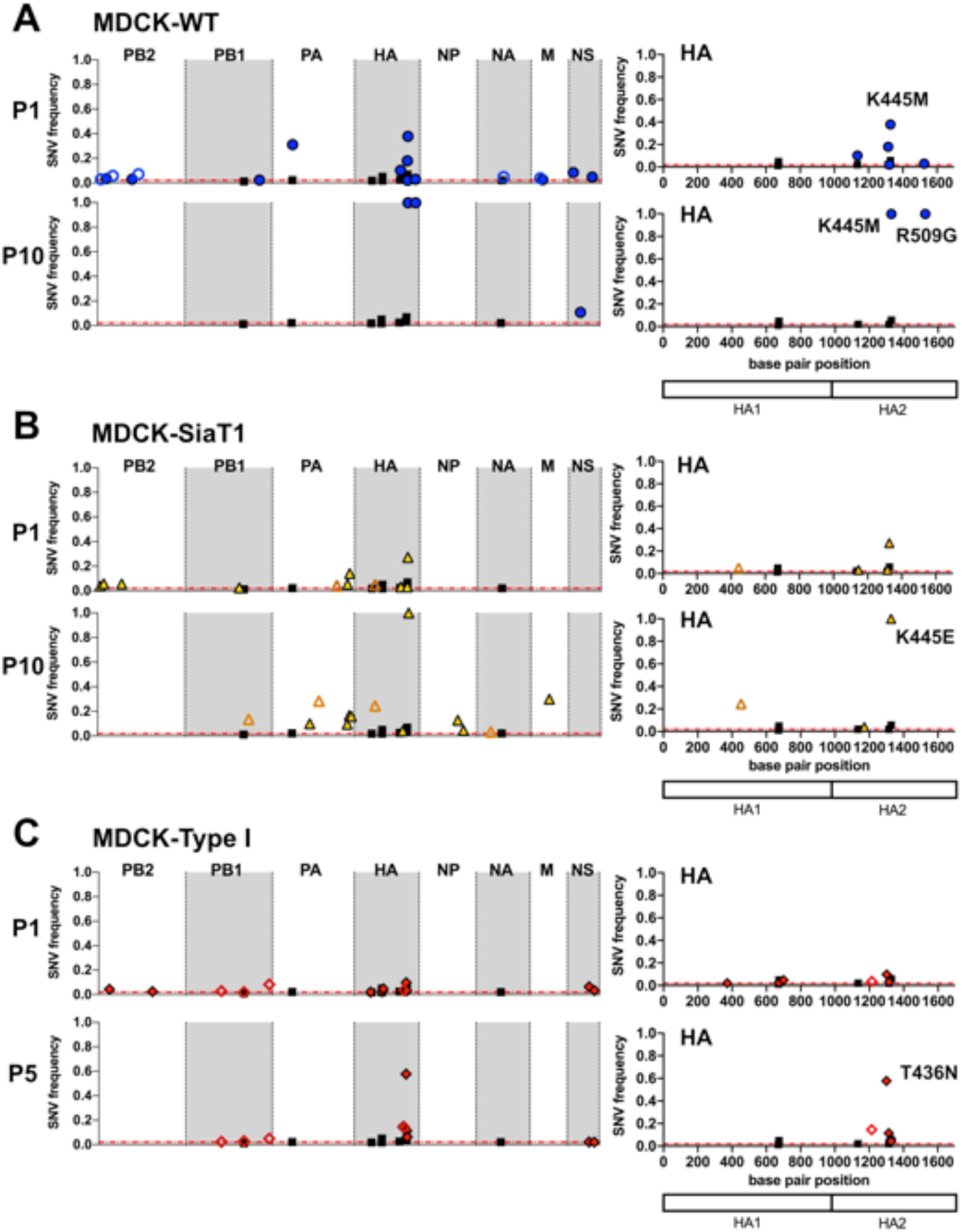
Genetic variation revealed in the pH1N1 virus. Single nucleotide variants (SNV) were analyzed during passage in **A)** MDCK-WT, **B)** MDCK-SiaT1, and **C)** MDCK-Type I cells. Data is shown for the whole genome for passage 1 and passage 10 (**A,B**) or passage 5 (**C**) on the left panel, while the nucleotide position of SNVs for HA are mapped on the right side with particular amino acid changes noted. Non-synonymous mutations are filled in shapes while synonymous mutations are open shapes.

**Figure 5.**
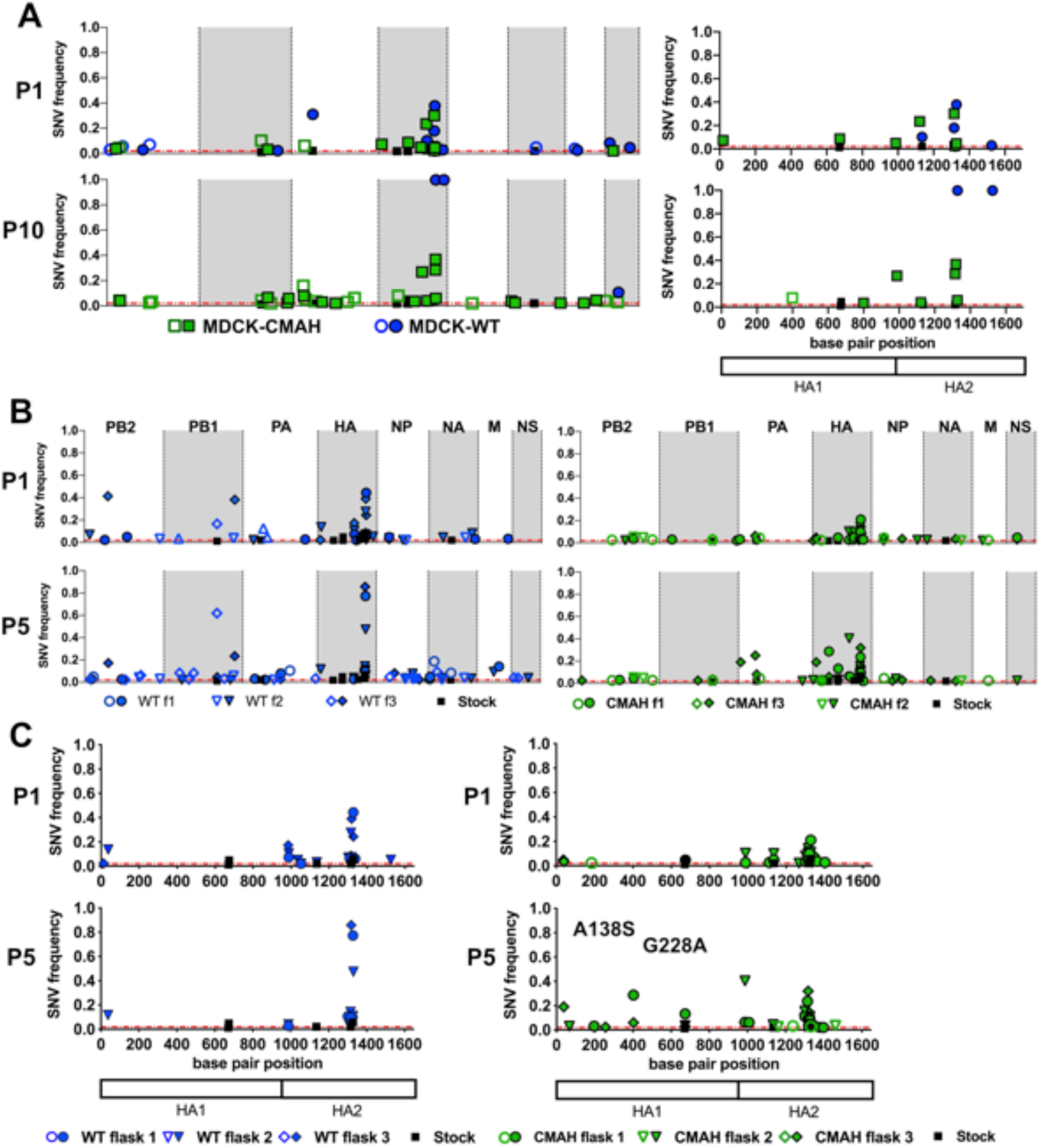
Single nucleotide variants (SNV) of pH1N1 passaged in MDCK-WT and MDCK-CMAH. **A)** SNVs from replicate 1 are mapped for the whole genome (left panel) and for HA alone (right panel) for passage 1 and passage 10. **B)** Whole genome map for SNVs from the three independent virus populations in replicate 2 of pH1N1 passaged in MDCK-WT or MDCK-CMAH. **C)** The nucleotide position of SNVs mapped for HA only from the three virus populations of pH1N1 in MDCK-WT or MDCK-CMAH. Particular amino acid changes as result of SNVs are also noted. Non-synonymous mutations are filled in shapes while synonymous mutations are open shapes.

### Passaging of pH1N1 in MDCK-CMAH cells

Initial passage of virus in MDCK-CMAH cells showed more nonsynonymous SNVs in the HA1 domain than in viruses passaged in the other cell types, so additional replicates of five passages in three separate virus populations were performed to determine the variation of the SNVs that arose (Fig. 2). Most SNVs in MDCK-CMAH passaged viruses were in the HA gene segment, along with changes in PA that were primarily synonymous and not consistent between virus populations (Fig. 5B). MDCK-CMAH passaged pH1N1 contained SNVs in the HA2 stalk domain at positions 441, 442, and 445, as was also seen in virus passed in other MDCK cell lines. Several apparently MDCK-CMAH specific SNVs arose near the receptor-binding site of HA during the later passages in these cells (Fig. 5**, Supplemental Table 2**). These include polymorphisms of residues 138 and 228 of HA1 in all MDCK-CMAH-passaged virus populations in both replicate 1 and 2, although these were not maintained through passage 10. Residues 138 and 228 fall within the outer loops of the Sia binding site and are known to alter α2,3- or α2,6-Sia binding preferences (42). Other SNVs near the receptor-binding site in MDCK-CMAH passaged virus populations showed variability (Fig. 5C), but none was present at >15% by passage 5, except for replicate 2 (A138S, ∼30%). After 10 passages in MDCK-CMAH cells, only one low frequency variant, I269N (4%), was present in HA1. Some pH1N1 virus passed in MDCK-CMAH cells also showed loss of proposed glycosylation sites 21 and 33 at low frequency (<10%) (Fig. 5B,C), similar to passages in MDCK-WT cells.

### Passage of wyoH3N2 in MDCK-WT, MDCK-SiaT1, and MDCK-Type I cells

The wyoH3N2 results varied, and the virus did not propagate past passage 3 in the MDCK-WT cells in the second replicate, likely due to the poor cell culture growth of recent H3N2 human viruses (43, 44). In other passage series in all MDCK cell lines, most SNVs were seen in the HA gene segment (Fig. 6A, B; **Supplemental Table 3**). Changes of residues in HA1 near the receptor-binding site of HA included residue N216K after 5 passages in both MDCK-WT and MDCK-Type I at 16% and 2% respectively (Fig. 6A,C), and in MDCK-WT N216K was >80% by passage 10. However, few SNVs in HA1 were seen in the MDCK-SiaT1 passaged virus and none were carried to passage 10, which is consistent with previous reports (Fig. 6B) (33).

**Figure 6.**
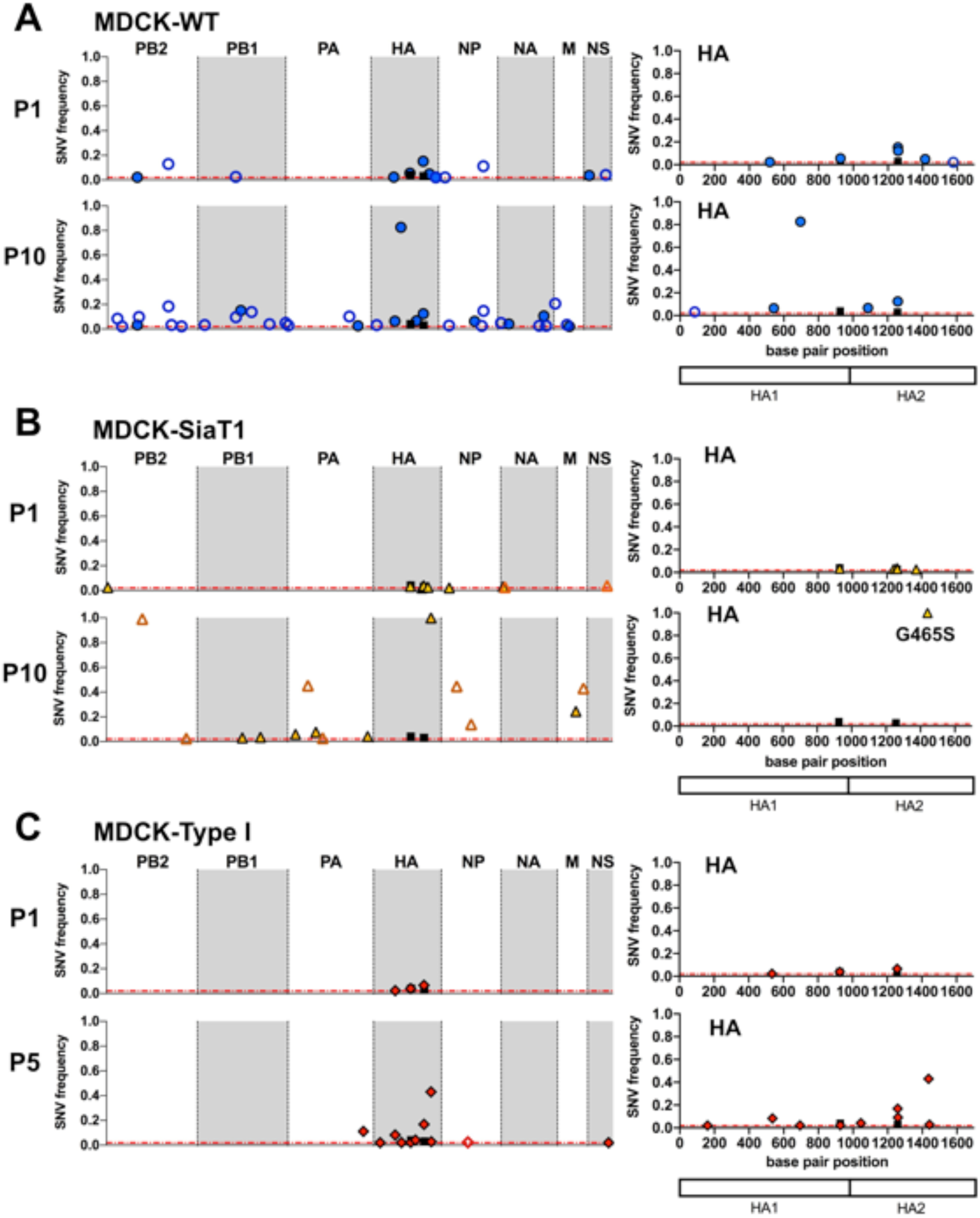
Genetic variation revealed in the wyoH3N2 virus. Single nucleotide variants (SNV) were analyzed for virus passaged in **A)** MDCK-WT, **B)** MDCK-SiaT1, and **C)** MDCK-Type I cells. Data is shown for the whole genome for passage 1 and passage 10 (**A,B**) or passage 5 (**C**) on the left panel, while the nucleotide position of SNVs for HA are mapped on the right side with particular amino acid changes noted. Non-synonymous mutations are filled in shapes while synonymous mutations are open shapes.

WyoH3N2 virus passaged in MDCK-WT also showed a low frequency loss of a glycosylation site at residue 165 in HA1 (N165T), which was maintained at a frequency of 5-10% through passage 10. SNVs in the stalk domain of HA2 occurred after passage in MDCK-WT, MDCK-Type I, and MDCK-SiaT1 cells (Fig. 6; **Supplemental Table 3**). HA residue 404 had two variants (G404R and G404E) in most populations at high frequency in passage 5 (16-75%) and G404E was maintained through passage 10 in MDCK-WT passaged virus at ∼12% frequency (Fig. 6A). Both variants were present in the stock virus at low frequencies (Supplemental Fig. 1). Other stalk mutations include G465S, which was fixed in MDCK-SiaT1 cells in passage 5 and maintained through passage 10 (Fig. 6B), and G463D at >40% in MDCK-Type I cells (Fig. 6C). Neither mutation was detected in the stock virus or the MDCK-WT grown virus. For the other gene segments, only a few low frequency (<3%) SNVs occurred, but most were synonymous mutations and not retained across passages (Fig. 6).

### Passage of wyoH3N2 in MDCK-CMAH cells

WyoH3N2 passaged in MDCK-CMAH showed more low frequency SNVs near the receptor-binding site than in other MDCK cell lines, although results varied between virus populations (Fig 7**; Supplemental Table 3**). SNVs included amino acid mutations of HA residues V186I, P221S, and Y233H at low frequencies (<5%) in both replicate 1 and replicate 2. WyoH3N2 passaged in MDCK-CMAH saw a high frequency SNV at residue N216K by passage 10 in replicate 1 (92%) similar to MDCK-WT passaged virus, as well as SNVs G404R/E (20-50% by passage 5), G465S (∼10% by passage 5), and the loss of the glycosylation site at position 165 (2-16%) that were also seen in other virus populations. However, only the D487N variant was maintained through all 10 passages in replicate 1. A small number of non-synonymous SNVs arose in the NA gene segment that led to changes in the proposed secondary Sia binding site of NA in MDCK-WT (S367G, ∼11%) and MDCK-CMAH (R428K, ∼37%). As this secondary Sia binding site has only recently been described (45), its importance for infection of cell cultures requires additional research. Additionally, the SNV F600S appeared repeatedly in the PA gene segment in MDCK-WT (∼42% in passage 5, lost by passage 10) and in MDCK-CMAH (∼10% in passage 5, ∼5% in passage 10) in replicate 1. This variant is located in the C-domain of PA that interacts with PB1; however, position 600 is not directly involved in this interaction and it is unclear if this variant has any function (46).

**Figure 7.**
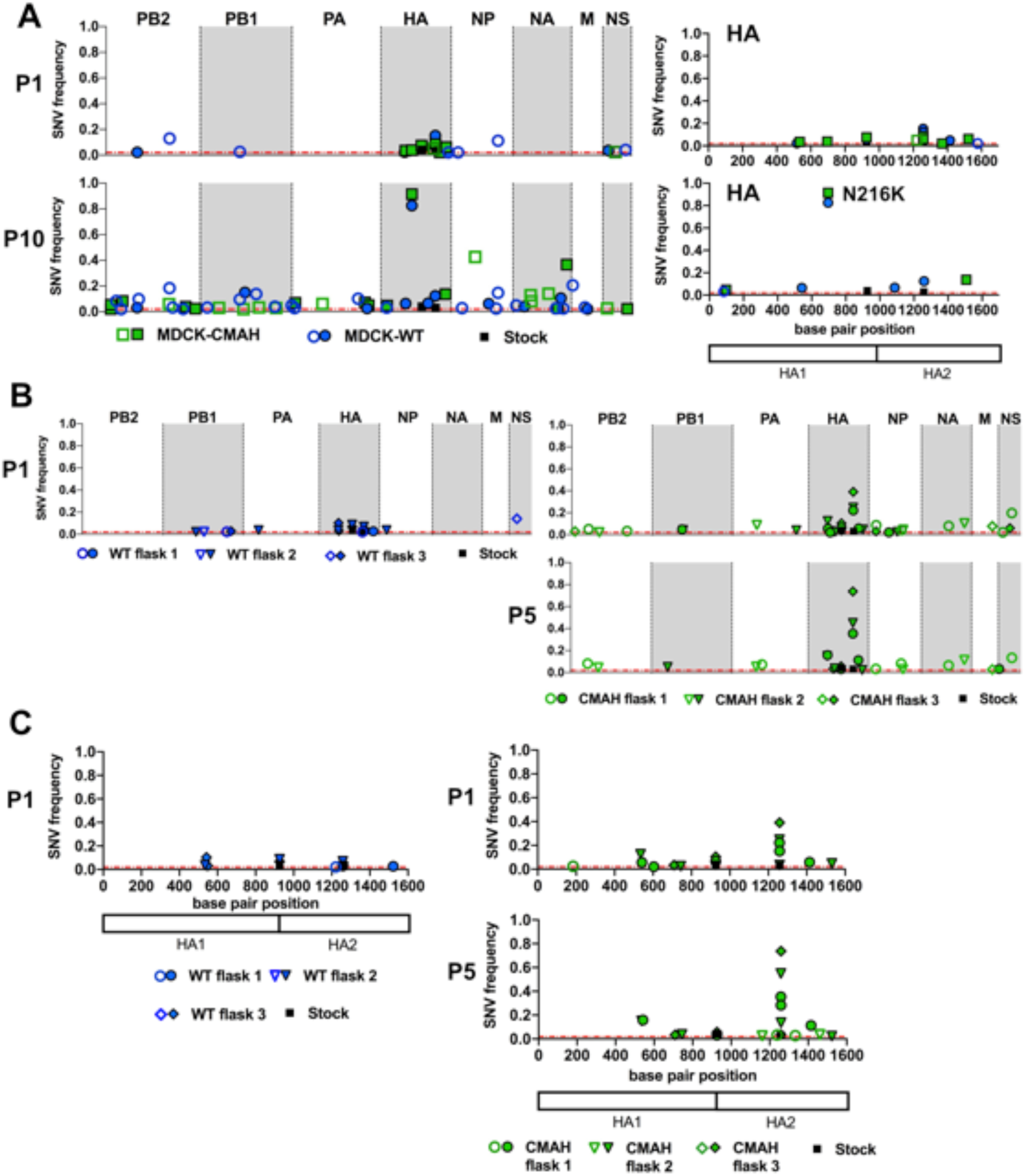
Single nucleotide variants (SNV) are mapped for wyoH3N2 passaged in MDCK-WT and MDCK-CMAH. **A)** SNVs from replicate 1 are mapped for the whole genome (left panel) and for HA alone (right panel) for passage 1 and passage 10. **B)** Whole genome map for SNVs from the three independent virus populations in replicate 2 of wyoH3N2 passaged in MDCK-WT or MDCK-CMAH. **C)** The nucleotide position of SNVs mapped for HA only from the three virus populations of wyoeH3N2 in MDCK-WT or MDCK-CMAH. Non-synonymous mutations are filled in shapes while synonymous mutations are open shapes.

### CIV H3N2 in MDCK-WT, MDCK-SiaT1, and MDCK-Type I cells

The CIV H3N2 showed little consistent HA stalk mutations, only comprising a SNV at position I335K that was also present in the stock virus and maintained at low frequency (Supplemental Fig. 1; Fig. 8). This variant was positioned just after the cleavage site between HA1 and HA2 and may be associated with protease susceptibility, although it was not maintained through longer passages. Other SNVs in HA varied between the replicates in MDCK-WT cells. Residue S186I was >50% in passage 5 and >90% by passage 10 in replicate 1, while the three populations in replicate 2 showed only low frequency SNVs in HA by passage 5 (Fig. 8A, Fig. 9B). Viruses grown in MDCK-Type I and MDCK-WT cells both showed SNVs at position 186 (∼50%) (Fig. 8C). In MDCK-SiaT1 cells, the SNV at position S219P in the receptor-binding site reached a frequency of 98% by passage 10 (Fig. 8B). Residue 219 may be associated with changes in receptor binding preference of human H3N2 virus passaged in eggs (47), and may be selected by α2,3-linked Sia, which is preferred by canine IAV strains, to the α2,6-linked Sia expressed in MDCK-SiaT1 cells.

**Figure 8.**
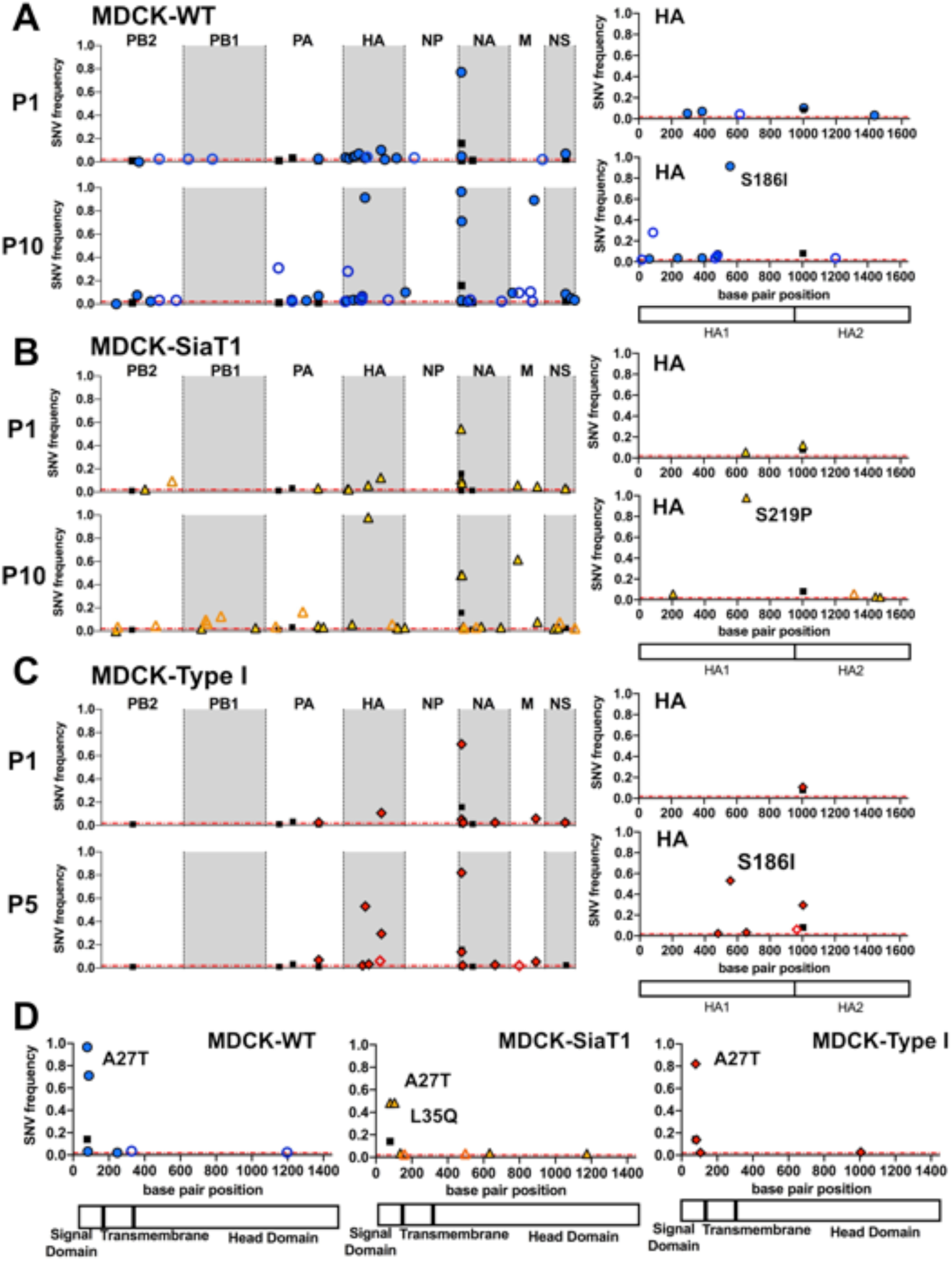
Genetic variation revealed in the CIV H3N2 virus. Single nucleotide variants (SNV) were analyzed for virus passaged in **A)** MDCK-WT, **B)** MDCK-SiaT1, and **C)** MDCK-Type I cells. Data is shown for the whole genome for passage 1 and passage 10 (**A,B**) or passage 5 (**C**) on the left panel, while the nucleotide position of SNVs for HA are mapped on the right side with particular amino acid changes noted. **D**) SNVs mapped for NA for passage 10 for MDCK-WT and MDCK-SiaT1 or passage 5 for MDCK-Type I. Non-synonymous mutations are filled in shapes while synonymous mutations are open shapes.

**Figure 9.**
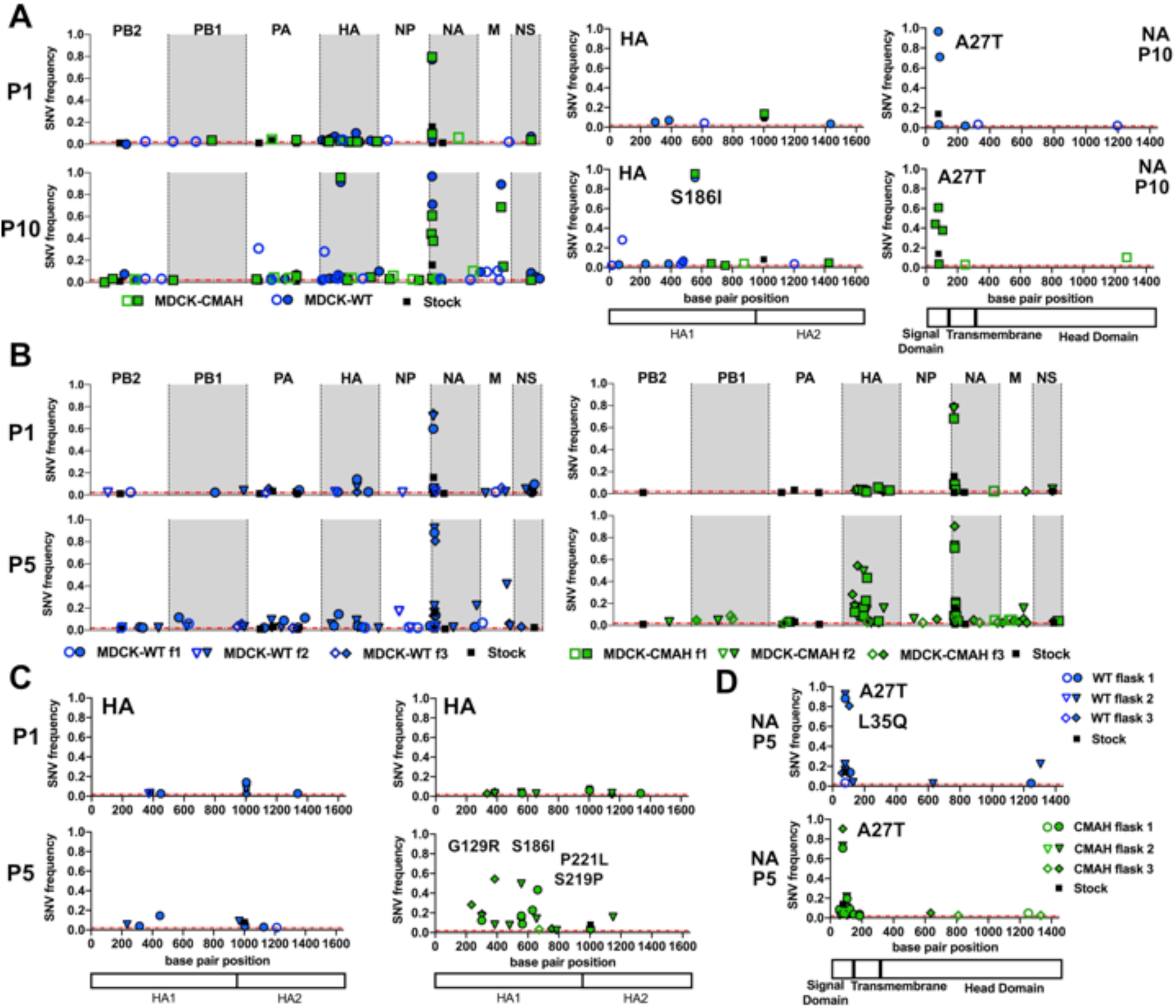
Single nucleotide variants (SNV) are mapped for CIV H3N2 passaged in MDCK-WT and MDCK-CMAH. **A)** SNVs from replicate 1 are mapped for the whole genome (left panels) and for HA alone (middle panels) and NA alone (right panels) for passage 1 and passage 10. **B)** Whole genome map for SNVs from the three independent virus populations in replicate 2 of CIV H3N2 passaged in MDCK-WT or MDCK-CMAH. **C)** Nucleotide position of SNVs mapped for HA only from the three virus populations of CIV H3N2 in MDCK-WT or MDCK-CMAH. **D)** Nucleotide position of SNVs mapped for NA only for the three virus populations of CIV H3N2 in MDCK-WT or MDCK-CMAH. Non-synonymous mutations are filled in shapes while synonymous mutations are open shapes.

An A27T variant in the NA was present in the stock virus (∼14%) and reached 60-95% by passage 5 in all cells (Fig. 8D). In passage 10 virus from MDCK-WT cells, A27T reached 97% frequency, while passage 10 virus from MDCK-SiaT1 contained both A27T (∼48%) and L35Q (∼48%). One MDCK-WT passage 5 virus population in replicate 2 also had the L35Q variant (∼80%) (Fig. 9D**; Supplemental Table 4**). Several additional SNVs occurred in the NA transmembrane and stalk domain by passage 5 and passage 10, including residues 28, 30, 35, 36, and 37 at frequencies of 5-70%. Similar mutations were also seen in canine H3N2 viruses when they were passaged on feline cells, so they may result from general cell culture adaptation for increased stability or budding efficiency (48).

The other gene segments of CIV H3N2 virus showed few consistent minority SNVs between passages and cell types. The C489S variant in PA was present in the stock virus and was maintained through passage 5 and passage 10 of all virus populations at <10% frequency. An NS segment variation within the NEP protein at position 35 was seen in all virus populations at low (<5%) frequency in passage 1 and was maintained in MDCK-WT grown virus through passage 10 at low frequency. Also, a G228R variant in M1 reach high frequency in MDCK-WT passaged virus by passage 10 and was also fixed in virus passaged with Zanamivir (Fig. 3C).

### CIV H3N2 in MDCK-CMAH cells

Viruses grown in MDCK-WT and MDCK-CMAH cells showed variation between replicates 1 and 2 virus populations (Fig 9A,B). However, more SNVs in HA1 arose in replicate 2 MDCK-CMAH passed virus compared to that in MDCK-WT cells, with variants near the receptor-binding site at residues 183, 186, 188, 219, and 221, some reaching frequencies of 20-70% (Fig. 9B,C; **Supplemental Table 4**). However, in replicate 1 only the S186I variant reached high frequency (∼96% by passage 10) in MDCK-CMAH passed virus, with other minority variants at positions 129, 183, 221, and 252 present at ∼10% or lower frequencies. The S186I was also reached high frequency (91%) in MDCK-WT passaged CIV H3N2 in replicate 1 by passage 10 (Fig. 9A). This mutation appeared in all MDCK-CMAH passaged viruses at frequencies of 17% to 67% by passage 5 in both replicates, but was present only in replicate 1 in MDCK-WT. The variants S219P and P221L are within the 220-loop of the receptor-binding site and were seen at low frequencies in several of the MDCK-CMAH populations by passage 5. P221L only occurred in MDCK-CMAH passaged virus (43% frequency in passage 5, replicate 2) while S219P also occurred in MDCK-SiaT1 passaged virus (98% by passage 10). Viruses passaged in MDCK-CMAH also showed variation of HA residue 335 in the stalk at low frequency, as well as SNVs in NA at positions 20, 27, 28, 35, and 36, with A27T having high frequency (70-90%) in all passage 5 virus populations (**Supplemental Table 4**). The low frequency C489S variant in PA, F35L in NEP sequence of the NS segment, and G228R variant in M were also seen in the MDCK-CMAH passaged virus.

### MDCK-Type II cells were less permissive to IAV infection

All three viruses replicated well in the MDCK-WT and MDCK-Type I cells, but showed lower replication in MDCK-Type II cells, even though all cell types expressed similar amounts of Sia on their surface (Fig. 1A). Lower numbers of infected cells were seen at early time points after infection (Fig. 10A) and significantly less virus was collected in the supernatant after 48 hours (Fig. 10B). Virus infections in Type-II cells were very sensitive to the presence of trypsin in the media, as had previously been reported (17). Only fresh stocks of trypsin added to infection media allowed for efficient infection. A higher MOI of 0.05 was used for passaging experiments due to the lower replication to assure high enough titers for RNA extraction and sequencing. SNVs primarily arose in the HA segment of the MDCK Type-II cells for all three viruses (Fig. 10C-E). However, most SNVs were low frequency, and those few SNVs that reached higher frequencies were also seen after passage on other MDCK cells (**Supplemental Tables 2, 3, 4**). Interestingly, CIV H3N2 did not replicate through all 5 passages, likely due to the lower infection efficiency of this strain compared to the human strains.

**Figure 10.**
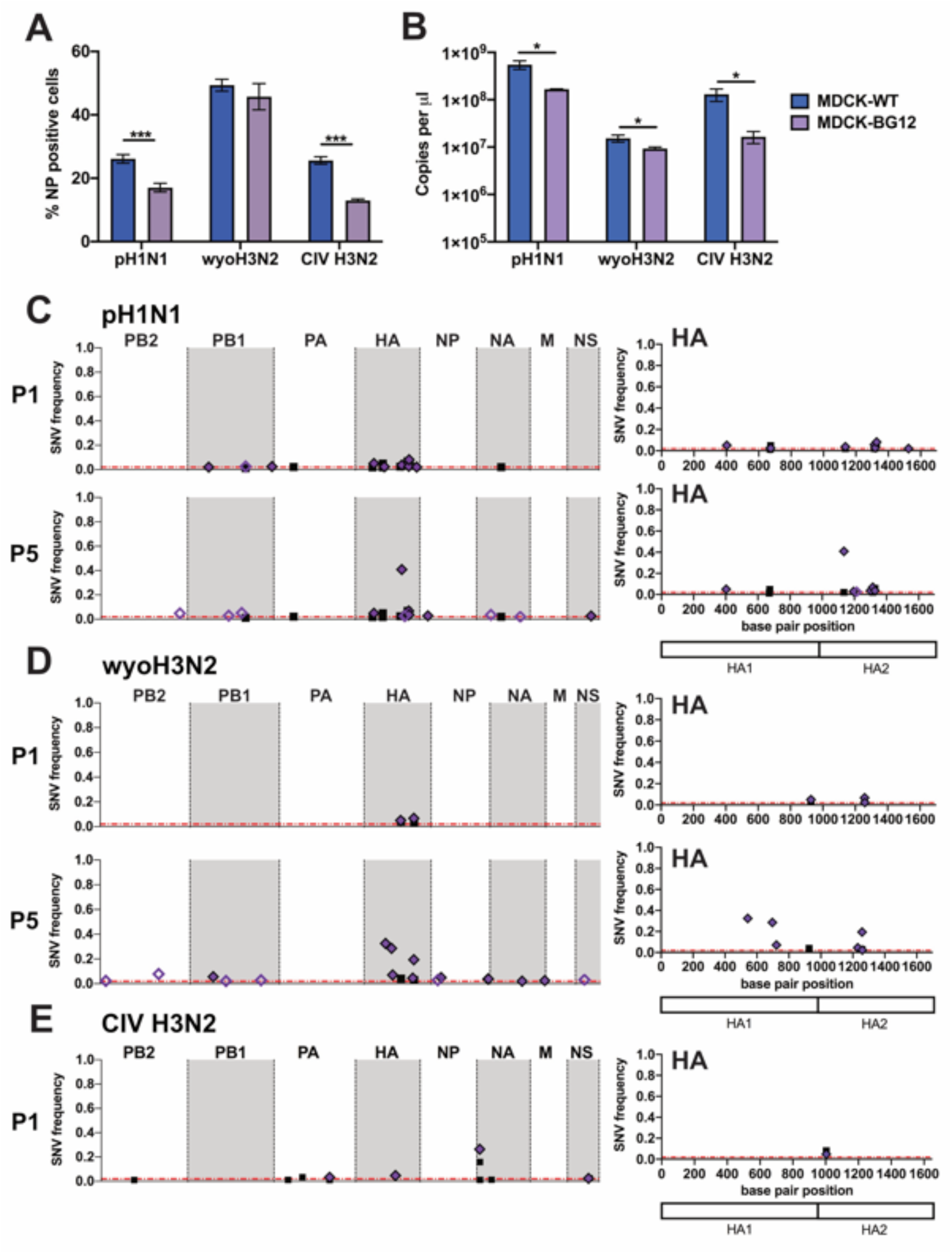
Analysis of MDCK Type-II cells, and results of viral passages. MDCK-Type II cells showed lower number of infected cells and lower levels of progeny virus than MDCK-WT cells. **A)** Flow cytometry of MDCK-WT and MDCK-Type II cells infected at an MOI of 0.25 for 6 hours, fixed and stained for NP protein. **B)** Supernatant from MDCK-WT and MDCK-Type II cells infected at an MOI of 0.01 for 48 hours was collected and virus titers determined by RT-qPCR for the M gene segment. Data analyzed by t-test using PRISM software. Single nucleotide variants (SNV) were analyzed for **C)** pH1N1, **D)** wyoH3N2, and **E)** CIV H3N2 passaged on MDCK-Type II cells. Data is shown for the whole genome for passage 1 and passage 5 on the left panel, while the nucleotide position of SNVs for HA are mapped on the right side. Non-synonymous mutations are filled in shapes while synonymous mutations are open shapes. For CIV H3N2, virus failed to replicate through passage 5, so only data for passage 1 is shown. **\*** = p-value ≤0.05; ** = p-value ≤0.01; *** = p-value ≤0.001.

## DISCUSSION

### MDCK cell types varied in virus infection efficiency

Many MDCK cell stocks, including the standard ATCC lineage, are known to be heterogeneous and to give rise to phenotypically distinct cells when passaged only 20 or more times, and some lineages differ in susceptibility to IAV strains (6, 17). While the MDCK-WT cells and MDCK-Type I cells examined here were quite susceptible and grew the virus to high titers, the MDCK Type II clone cells were infected at much lower levels by all three IAVs tested. A previous study with similar cells showed that some clones lacked necessary levels of proteases to activate HA for infection (17). We found that fresh trypsin stocks were necessary for efficient infection, likely due to this increased dependence on protease in the infection media. It is not clear whether the other differences between the Type I and Type II cells influence their ability to be infected or replicate virus, such as their glycolipid composition, metabolism, or polarization (13–15). It is also noteworthy that the CIV H3N2 virus had the lowest infection efficiency in MDCK-Type II cells and was not maintained through the 5 passages of our study. The cause of this difference is unknown as the canine virus should not be disadvantaged growing in canine cells, although of kidney epithelial origin.

### Only low levels of variation were generally detected in viruses passaged in different MDCK lineages

The three virus stocks used here all derived from reverse genetics plasmids and were prepared in MDCK-WT cells. Few variants were detected in the starting virus populations cells and they were primarily in the HA segment. After repeated passages of the viruses in the different MDCK cell lines, few SNVs were present within most of the viral gene segments, with most again present in the HA gene segment (Fig. 11), and in the NA segment for CIV H3N2. Using a threshold of 2% allowed us to focus on mutations that were more likely to be biologically significant. While replication in cell culture lacks many of the selective pressures found in natural infection, low levels of diversity within the IAV sequences in natural samples have been reported in human, equine, and canine infections (49–52). Similar results were seen here, with similarly low levels of variation in the sequences of these same strains of IAV passaged in wild type or CMAH^-/-^ mice (38). Some minority variants arose in individual virus populations in our different replicate experiments, in which three virus populations were passaged 5 times in MDCK-WT or MDCK-CMAH cells, revealing little variation within populations, but some variation - likely stochastic - between populations. This would align with the expectation that intra-host variation would be much lower compared to the variation seen between IAVs circulating in humans or other hosts, where additional sources of variation would include genetic drift, antigenic selection, and bottlenecks of varying sizes over extended chains of transmission (49).

**Figure 11.**
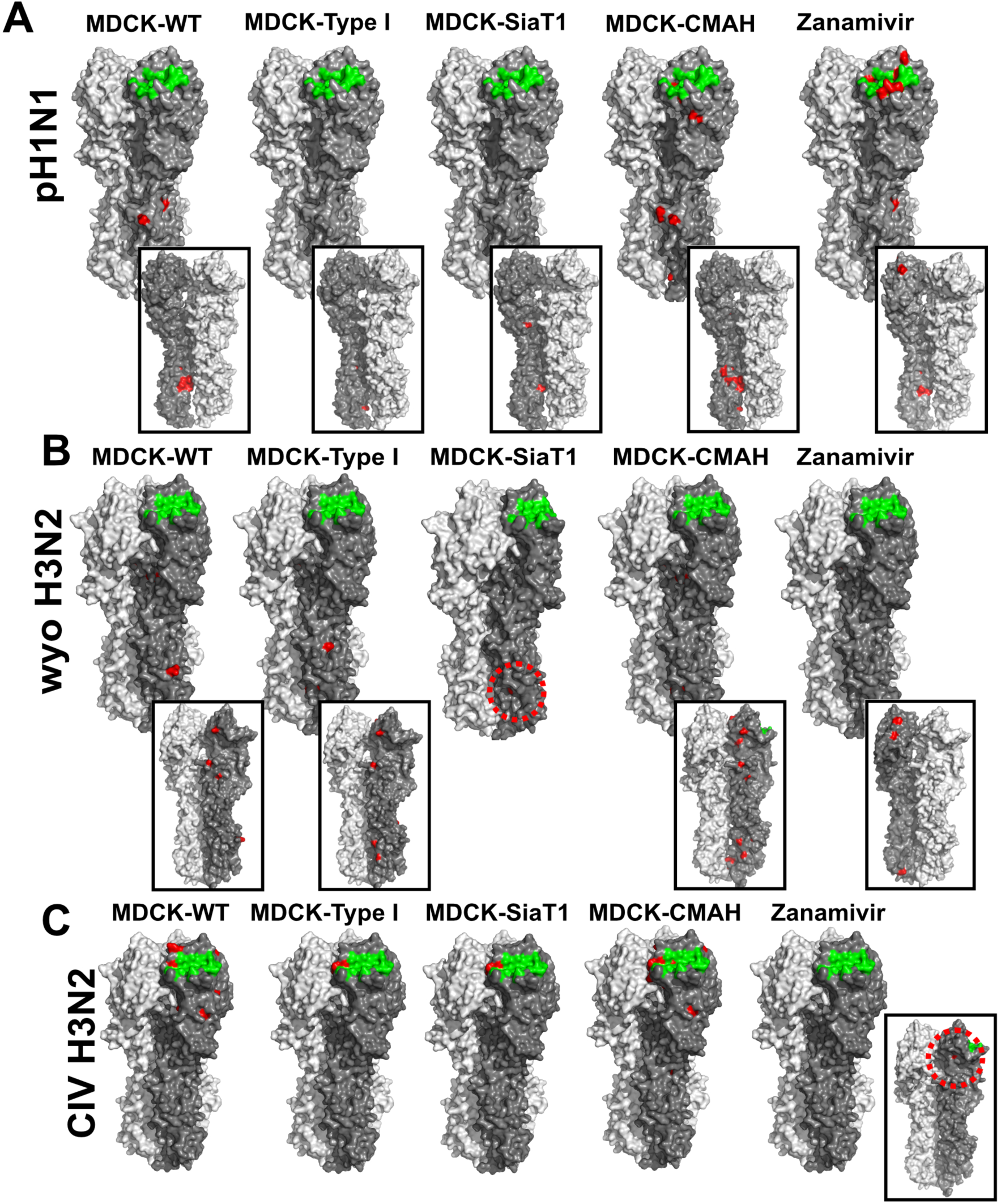
Many mutations seen occurred in the HA protein. Models of IAV HA with receptor binding (green) and amino acid changes (red) for each passage virus population. Inset images show the interior of the HA trimer with one monomer removed to highlight amino acid variants that are not solvent exposed. HA models were created using PYMOL with structures from A/Aichi/2/1968 H3 (2YPG) and A/California/04/2009 H1 (3LZG).

### Mutations in the HA gene were more frequent

For both pH1N1 and wyoH3N2 human IAV strains, most amino acid mutations altered residues within the stalk region of the HA gene, with some increasing to high frequency between 40% and 99%. Similar mutations altering the pH stability of the HA influence host tropism and stability during transmission (53, 54). These mutations may therefore be due to selection for HA stability at different pHs, or for more efficient protease cleavage or fusion in MDCK cells under the culture conditions of our studies. Comparing the variant sequences to those in databases showed that many of the stalk mutations that reached fixation here are present in other IAV isolates. In pH1N1, the K445E/M mutations were also present in some swine viruses and a few human isolates present on NCBI/Genbank. However, where culture information was available, most isolates containing these mutations had been passed in MDCK cells, so it is possible that these also represent culture adaptation. The G404R/E mutations that arose in several wyoH3N2 populations, however, were primarily seen in equine and canine IAVs, but the lack of information about passaging history of many of the isolates makes it difficult to draw any strong conclusions. SNVs in the HA stalk were not as prevalent in CIV H3N2 virus, so are likely to be associated with the specific sequences of the human viruses, or their replication in MDCK cells.

### Sialic acid linkages and effects on viruses

Human IAV favor cell binding and infection using the α2,6-linked Sia which is common in the human upper respiratory tract. The MDCK-SiaT1 cells have higher expression of this form of Sia linkage. Human viruses passaged in wild-type MDCK cells often have mutations around their Sia binding sites (32, 55). We saw no significant difference in the SNVs near the HA receptor-binding sites of pH1N1 when passed in MDCK-WT or MDCK-SiaT1 cells. In wyoH3N2, a mutation at position 216 near the receptor-binding site reached a high frequency when passed in MDCK-WT but no mutations in HA1 arose in MDCK-SiaT1 cells, confirming that increased α2,6-linked Sia maintains virus receptor-binding preference. In our study, this appears to be more important for human H3N2 than H1N1 viruses. CIV H3N2 virus specifically binds α2,3-linked Sia (56), and virus passaged in MDCK-SiaT1 cells showed near fixation of residue S219P near the receptor-binding site, suggesting selection by the Sia linkage type. CIV H3N2 virus also showed a number of SNVs after passage in various MDCK cell lines, mostly in the NA gene close to or within the trans-membrane and stalk region, centered around position 27. Some of these mutations rose to near fixation in all CIV H3N2 passages, suggesting that these mutations increase stability or budding efficiency in cell culture. Similar mutations have been reported in canine IAV strains passaged in feline cells (48). While some of these mutations did not correspond to any IAV isolates in the NCBI database (A27T and L35Q), others were seen in natural canine and avian isolates.

### Cell expression of Neu5Gc shows varying effects

Most IAV that have been tested preferentially bind Neu5Ac, the principle Sia form found in humans, birds, ferrets, Western breeds of dogs, and on MDCK cells (57, 58). However, pigs and horses do express high levels of Neu5Gc in their respiratory tracts (26, 27). Some viruses from horses, particularly the H7N7 strain that circulated from 1956 until the mid-1970s, appear to bind Neu5Gc preferentially (28). Laboratory-generated HA proteins that bind Neu5Gc had a T155Y mutation, with residues 143 and 158 also being related to this shift in Sia binding preference (59). Passaging of all three IAVs in our MDCK-CMAH cells, which express ∼40% Neu5Gc, resulted in more low level SNVs near the receptor-binding site compared to viruses passaged in MDCK-WT cells, but the mutations did not match the previously described mutations that changed binding from Neu5Ac to Neu5Gc (28, 59). These low frequency SNVs in the HA1 region that arose in MDCK-CMAH passaged virus populations were similar to mutations affecting α2,3- or α2,6-linked Sia binding (42). Other mutations in MDCK-CMAH passaged virus fell within other regions of the HA head domain (60). As Neu5Gc only comprised ∼40% of Sia on the cells, viruses may have been using the remaining Neu5Ac on the cell surface. However, the smaller pool of available receptor may have still resulted in selection for mutations in or near the receptor-binding site (28). The levels of Neu5Gc in our MDCK-CMAH cells were similar to those found in pig respiratory tract, where they may affect receptor-binding preferences and alter antigenic sites during natural infection in pigs.

### Passage in the presence of Zanamivir led to mutations in HA across all three viruses

As a control for the emergence of mutations under direct selection, we also passaged each virus with increasing concentrations of 2-deoxy-2,3-didehydro-*N*-acetylneuraminic acid (DANA or Zanamivir). By passage 10, pH1N1 did not show any SNVs in NA, but some SNVs did arise in HA near the receptor binding site, all of which remained <25% of the population. Two SNVs arose the in HA of WyoH3N2 near the receptor-binding site (positions 163 and 244), while a S231G mutation reached fixation in the HA of CIV H3N2. Previous studies have found that IAV strains can avoid Zanamivir inhibition through mutations in HA that change Sia binding affinity, so that the mutations we saw in HA could be associated with Zanamivir resistance (39, 41, 61, 62).

In summary, passage of the three viruses up to 10 passages in the variant MDCK cells or those expressing different Sia forms resulted in only a small number of changes in the HA gene segment, or a small number of variants in the NA gene segment of CIV H3N2, with little synonymous or nonsynonymous variation in the other gene segments. It can be difficult to distinguish whether a mutation is being directly selected or whether a mutation is neutral or physically linked to an advantageous mutation as it rises in frequency. While we assume that the mutations we saw arise in HA are being directly selected due to receptor variation or NA inhibition, further functional characterization will be needed to confirm this hypothesis. However, these experiments reveal how passage in different MDCK cell cultures or in the presence of different Sia or linkages may shape different IAV viruses. This is relevant for developing IAV vaccines in mammalian cells and for properly annotated wild isolates that are passaged in cells during characterization. Additionally, it confirms that diversity of the sequences of IAV populations are generally low, as previously seen in viruses in both *in vivo* experiments and clinical samples.

## MATERIALS AND METHODS

### Cells and viruses

MDCK-NBL2 and HEK293T cells were obtained from ATCC. MDCK-SiaT1 cells were prepared by transfection of the *ST6Gal1* gene in a plasmid under the control of the CMV promoter (pcDNA3.1, Invitrogen). Cell clones with increased levels of α2,6-linked Sia were identified by staining with the *Sambucus nigra* (SNA) lectin (Vector laboratories). MDCK-CMAH cells were prepared by transfection of the human CMAH gene in a plasmid under the control of the CMV promoter (pcDNA3.1, Invitrogen). Clones with Neu5Gc expression were determined by HPLC analysis as previously described (63). MDCK-Type I cells (clone AA7) and MDCK-Type II cells (clone BG12) were a gift from Dr. William Young (University of Kentucky) and were originally cloned and characterized by Dr. Guy E. Nichols (11). All cells were grown in DMEM with 10% fetal calf serum, and 50µg/ml gentamycin.

Three IAV strains were derived from reverse genetics plasmids, comprising (i) human H1N1 pandemic IAV (A/California/04/2009, pH1N1) in plasmid pDP2002, (ii) human H3N2 seasonal IAV (A/Wyoming/3/2003, wyoH3N2) in plasmid pDZ, and (iii) a canine H3N2 IAV (A/Canine/IL/11613/2015, CIV H3N2) in pDZ. The plasmid encoding each viral segment was prepared from a single bacterial colony, and an 8-plasmid mixture for each virus was prepared and used for transfection of a 3:1 co-culture of HEK293T cells and MDCK cells (MDCK-SiaT1 cells for wyoH3N2). Each virus was passaged two additional times in the same MDCK, or MDCK-SiaT1 cells, to generate a passage-3 stock, which was tested for infectivity by TCID_50_ assay. Each plasmid mixture (as DNA) and the virus stocks were then used to generate libraries for Illumina sequencing, as described below, revealing the original sequences and any baseline variation of the virus populations used for cell passaging.

### Genotyping of MDCK cells

DNA extracted from the three cell lines was genotyped on the 220k semi-custom CanineHD array (Illumina, San Diego, CA). Genotype data were managed in PLINK 1.9 (www.cog-genomics.org/plink/1.9/) (64, 65). Single nucleotide polymorphisms (SNPs) with ≥95% missingness were removed, sexes were assigned based on Y chromosome SNPs and checked with X chr SNPs (using PLINK’s --check-sex command). After filtering, 214,325 variants remained for analysis. To look at relatedness between the cell line samples, an identity-by-descent (IBD) metric, pi-hat, was calculated in PLINK. Pi-hat is an estimate of the probability of sharing 0, 1 or 2 alleles between each pair of individuals.

### Lectin staining by flow cytometry

Cells were seeded at sub-confluency and incubated overnight at 37°C and 5% CO_2_. Cells were collected using Accutase (Sigma) to retain surface glycans, then fixed in 4% PFA for 15 min. Cells were blocked using Carbo-Free Blocking Solution (Vector Laboratories). Fluorescein-conjugated plant lectins from *Sambucus nigra* (SNA) and *Maackia amurensis* lectin I (MAA I) (Vector Laboratories) were diluted 1:1200 in blocking buffer and incubated with cells for one hour. A Millipore Guava EasyCyte Plus flow cyotometer (EMD Millipore, Billerica, MA) was used to collect data, and analysis was done using FlowJo software (TreeStar, Ashland, OR). Statistical analyses were performed in PRISM software (GraphPad, version 8).

### Passaging of virus in cells

For virus passaged in MDCK-WT, MDCK-SiaT1, MDCK-CMAH, and MDCK-Type I cells, virus was passaged at low MOI of 0.001 for passage one and two, then blind passaged for passage three through five or ten, depending on cell type (see Figure 3). For MDCK-Type II cells, an initial MOI of 0.05 was used for passage one and two followed by blind passaging for five total passages. For the Zanamivir passaged virus, Zanamivir concentration was 0.01μM for passage one and increased gradually to a final concentration of 50 μM by passage ten. Virus was inoculated at a constant MOI of 0.005 following titering after each passage. For infections, cells were seeded at sub-confluency in T12.5 flasks and allowed to settle for 4 to 6 hrs. Once settled, cells were washed and inoculated with virus diluted in infection media (DMEM with 0.3% BSA and 1μg/ml TPCK-treated trypsin (Sigma Aldrich)) and allowed to adsorb for one hour. Inoculum was removed and infection media replaced. Virus was collected at 48 hrs.

### Flow cytometry and titering of virus from MDCK-Type II cell infections

For analysis of infected cells via flow cytometry, cells were seeded into 12-well plates to be near confluency and allow to settle for 8 hrs before infection at a MOI of 0.25. At 6 hours post infection, cells were trypsinized and fixed with 4% paraformaldehyde and stained with a mouse anti-NP antibody and fluorescence-conjugated goat anti-mouse secondary. A Millipore Guava EasyCyte Plus flow cyotometer (EMD Millipore, Billerica, MA) was used to collect data, analysis using FlowJo software (TreeStar). Statistical analyses were performed in PRISM software (GraphPad, version 8). For titering of virus, cells were seeded in 12-well plates as described and infected at an MOI of 0.01 and supernatant was collected at 48 hrs post infection. Viral RNA was isolated using the QIAamp Viral RNA Mini Kit (QIAGEN). Influenza genome copies were quantified from RNA isolations by reverse transcription and quantitative PCR (RT-qPCR) for the M segment modified from CDC protocol (66). Products were amplified using Path-ID (Applied Biosystems) with M-specific primers (5’ to 3’, F: GACCRATCCTGTCACCTCTGAC, R: AGGGCATTYTGGACAAAKCGTCTA), probed with 5’-TGCAGTCCTCGCTCACTGGGCACG-3’ and run on a 7500 Fast Real-Time platform against a standard curve.

### Viral RNA extraction and virus titering

Viral RNA (vRNA) was isolated from cell culture supernatant using the QIAamp Viral RNA Mini Kit (QIAGEN). Influenza virus titers were determined by TCID_50_ on MDCK-NBL2 cells.

### Library generation and NGS sequencing

Library generation was performed as previously described (38). In brief, total vRNA was incubated with Superscript III and Platinum Taq-HiFi (Invitrogen) in the presence of universal influenza amplifying primers (5’ to 3’, uni12a: GTTACGCGCCAGCAAAAGCAGG; uni12b: GTTACGCGCCAGCGAAAGCAGG; uni13: GTTACGCGCCAGTAGAAACAAGG). For CIV H3N2, HA-specific primers (forward: AGCGAAAGCAGGGGATACT; reverse: CCAGTAGAAACAAGGGTGTTTT) were used in addition to the universal primer set to increase amplification of this segment. Viral cDNA products were purified by Agencourt AMPureXP magnetic beads (Beckman Coulter) and quantified by QuBit (Invitrogen). Sequencing libraries were prepared using Nextera XT DNA Library Prep Kit (Illumina). Pooled libraries were run on an Illumina MiSeq v2 for 250bp paired reads in the Cornell Genomics Facility of the Biotechnology Research Center.

### Sequence analysis and variant calling

Analysis was performed in Geneious v.11.1.5. Read trimming was performed by the BBDuk script (https://jgi.doe.gov/data-and-tools/bbtools/bb-tools-user-guide/bbduk-guide/), followed by read merging, and alignment to the reference genomic plasmid sequence for each virus. For variant calling, we considered those at >2% frequency with minimum 200 read coverage.

### Graphing and analysis

All figure graphs were generated in GraphPad Prism v.8.1.2.

### Data Availability

Sequences have been deposited at NCBI Sequence Read Archive (SRA) under BioProject PRJNA601451.

## ACKNOWLEDGMENTS

We thank Wendy Weichert for expert technical help and coordinating sequencing of canine cells. We also would like to thank Dr. Edward Holmes (University of Sydney) for his invaluable feedback and suggestions, and Dr. Henry Wan (University of Missouri) for the swine tissue samples.

---

**Supplemental Table 1.** HPLC analysis for total sialic acid content for tissue from pigs. Data given as percent of total Sia averaged across three runs.

| **Tissue** | **%Neu5Gc** | **%Neu5Ac** | **%Neu5,9Ac2** |
| --- | --- | --- | --- |
| Upper Lung | 36.8 | 51.0 | 0.7 |
| Lower Lung | 37.9 | 52.0 | 0.5 |
| Lower Trachea | 14.4 | 81.4 | 0.3 |
| Upper Trachea | 14.1 | 83.9 | 0.3 |

**Supplemental Figure 1.**
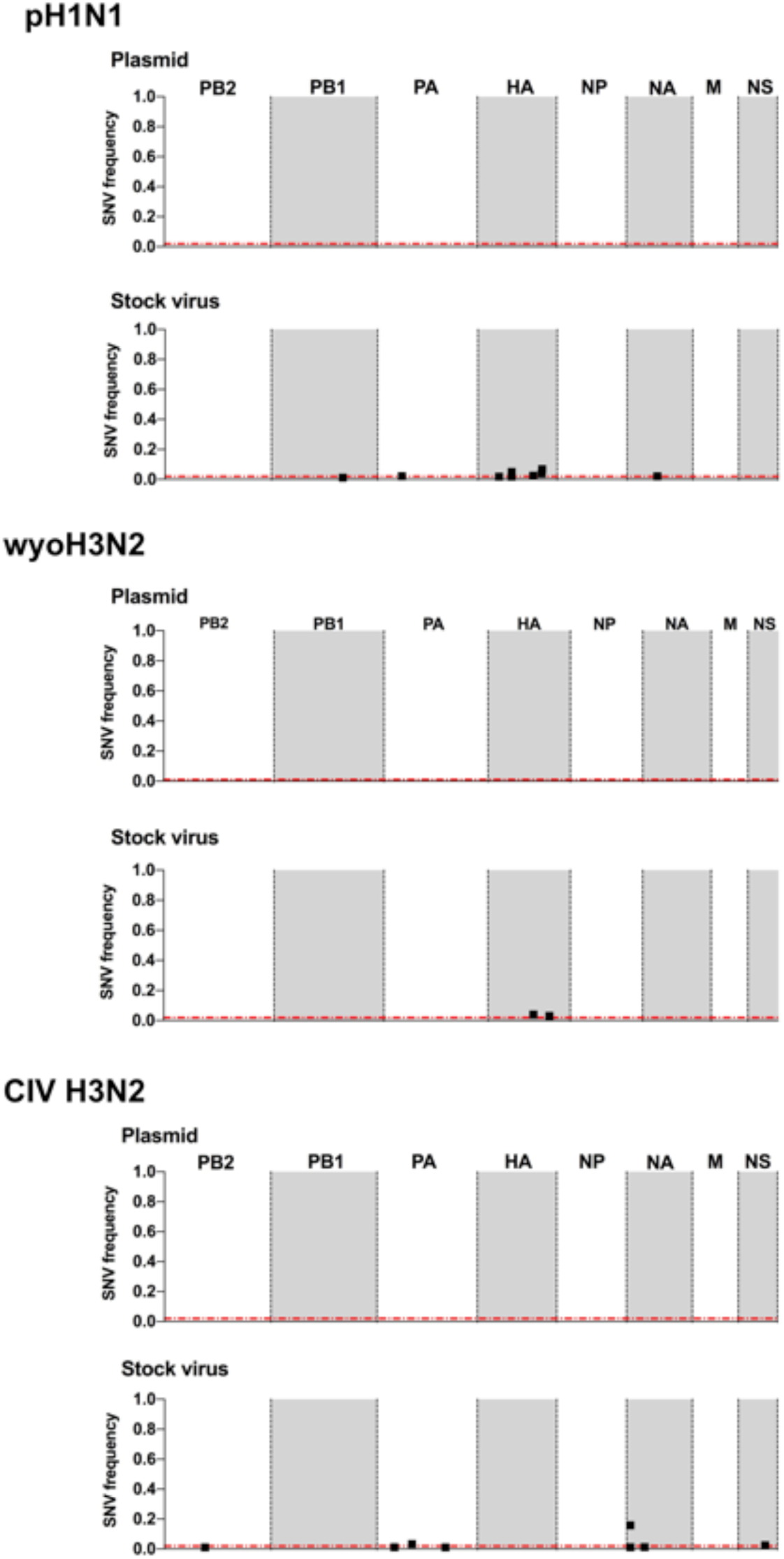
Results of complete genome deep sequencing studies, showing the single nucleotide variants (SNV) in plasmids and stock virus were analyzed for pH1N1, wyoH3N2, and CIV H3N2. Data is shown for the whole concatenated genome. Non-synonymous mutations are filled in shapes while synonymous mutations are open shapes.

**Supplemental Figure 2.**
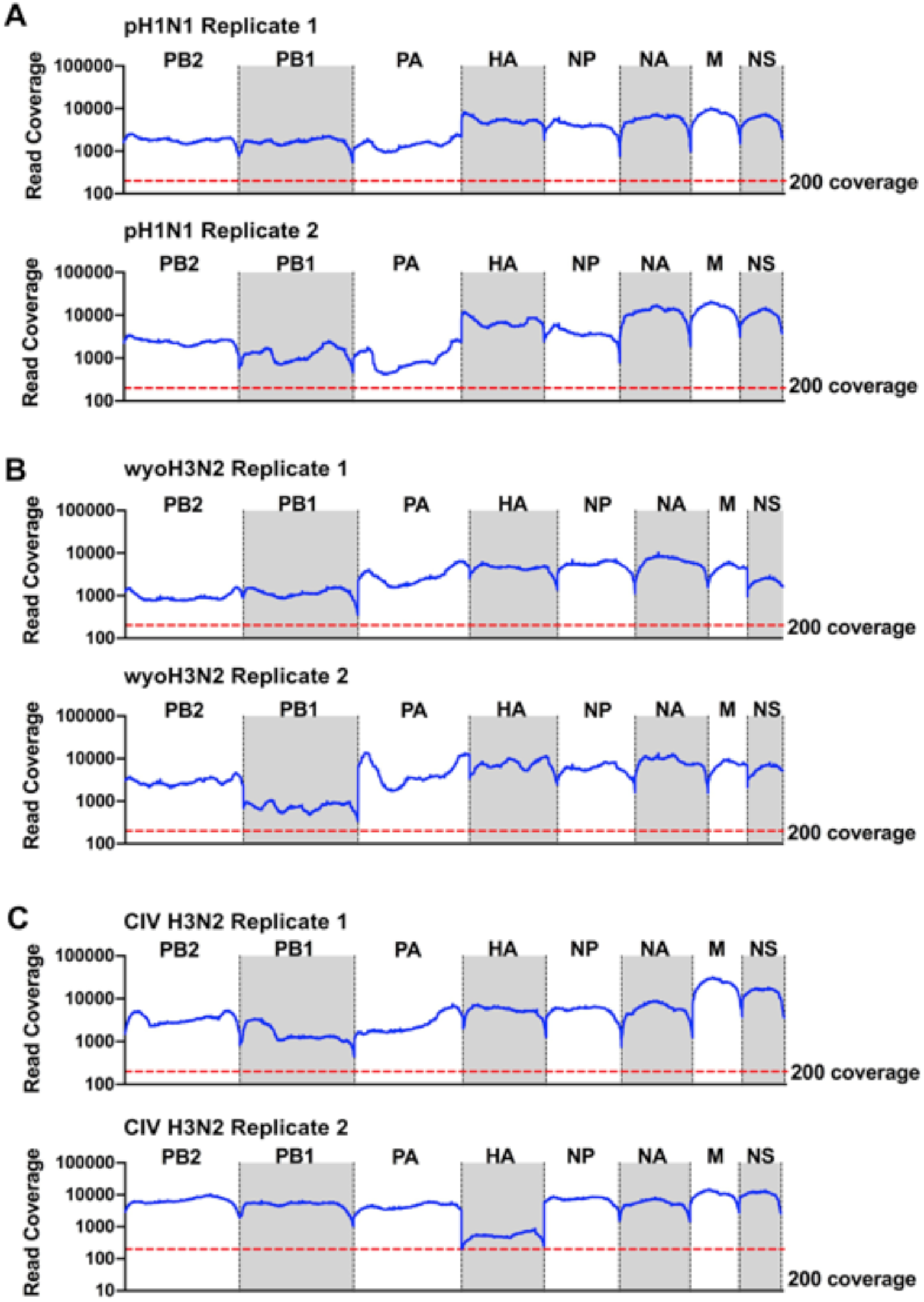
Average coverage for each genome segment for Replicate 1 and Replicate 2 for all viruses **A)** pH1N1, **B)** wyoH3N2, and **C)** CIV H3N2. For analysis, a cutoff of 200 reads per base pair was set.

**Supplemental Table 2. pH1N1 all mutations**

**Supplemental Table 3. wyoH3N2 all mutations**

**Supplemental Table 4. CIV H3N2 all mutations**

